# Integrated transcriptomic meta-analysis provides novel insights into breast cancer molecular subtypes

**DOI:** 10.1101/2021.08.04.455013

**Authors:** Akhilesh Kumar Bajpai, Sravanthi Davuluri, Kavitha Thirumurugan, Kshitish K. Acharya

## Abstract

Breast cancer is the most common cancer in women worldwide. There are four major breast cancer subtypes (luminal A, luminal B, HER2-enriched and triple-negative/TNBC). TNBC is the most aggressive form with the worst prognosis. However, the differences among the subtypes have not been completely established at the molecular level, thereby limiting therapeutic and diagnostic options for TNBC. We performed a meta-analysis of microarray and RNA-sequencing data to identify candidate genes with an expression-based association in each molecular subtype of breast cancer. The protein interaction network of the candidate genes was analyzed to discover functionally significant gene clusters and hub genes. Potential therapeutic candidates for TNBC were explored through gene-miRNA interactions. We identified 316, 347, 382, and 442 candidate genes in luminal A, luminal B, HER2 and TNBC subtypes, respectively. A total of 135 (26 up- and 109 down-regulated) candidate genes were shared by all four subtypes. Functional analysis of the candidate genes indicated up-regulation of ‘cell cycle’ and ‘p53 signaling’ pathways and down-regulation of multiple signaling pathways. COL10A1 was found to be highly up-regulated in all subtypes. It may be a good target for research towards multiple types of applications, including therapeutics. KIF4A, a commonly up-regulated X-chromosome gene was significantly associated with the survival of breast cancer patients. Protein interaction network and centrality analysis revealed that low-moderately differentially regulated genes play an important role in functional cascades across proteins in breast cancer subtypes and may be potential candidates for therapeutics. Targeting FN1 (fibronectin 1), the key up-regulated hub gene by miR-1271 5p may be an important molecular event to be targeted for potential therapeutic application in TNBC.

## Introduction

Breast cancer is the most common cancer in women, accounting for 24% of all the cancer cases, and with an estimated number of ~2 million cases worldwide. It is responsible for ~600,000 cancer related deaths and accounts for 15% of all cancer related mortalities in women worldwide (Torre et al. 2016). Based on the expression of signature genes, breast cancer has been classified into four major molecular subtypes: luminal A, luminal B, Her2-enriched (HER2), and basal-like/triple-negative breast cancer (TNBC) (Perou et al. 2000; Sorlie et al. 2001). Luminal A is the most common sub-type (~50% cases) followed by luminal B (~20% cases), HER2 and TNBC (~15% each). Among these, luminal A has the best prognosis and TNBC has the worst. TNBC is the most aggressive sub-type, which is responsible for poorer survival and higher incidences of distant metastasis (Anders and Carey 2009). There are limited targeted therapy options for TNBC due to the lack of expression of estrogen (ER), progesterone (PR), and HER2 receptors (Bianchini et al. 2016; Kumar et al. 2017).

High-throughput technologies, such as microarray and RNA-sequencing have been used extensively to study the mechanisms of various diseases including breast cancer. Multiple studies have used these large-scale datasets to identify better prognostic and therapeutic markers in breast cancer subtypes, understand the underlying disease mechanisms, identify novel metastatic genes, and predict metastatic outcomes. With the availability of multi-omics high-throughput data, it has been possible to explore the cellular systems as a network of interacting molecules, including proteins, miRNAs, and lncRNAs etc. For instance, a study by Hsu et al., (Hsu et al. 2019) identified six novel immunoglobulin genes as biomarkers for better prognosis of triple-negative breast cancer using publicly available microarray studies. Similarly, Dong et al., (Dong et al. 2018) identified key genes and pathways in triple-negative breast cancer based on an integrated bioinformatics approach. A study by Guo et al., (Guo et al. 2017) identified and validated nine novel TNBC associated biomarkers by integrating gene expression data and protein interactions. Furthermore, microarray gene expression data was used to discover novel genes associated with lymph-node metastasis in TNBC subtype (Mathe et al. 2015). Also, the authors correlated the gene expression changes with miRNA and confirmed the relations hip between specific miRNAs and lymph-node metastasis. Yu et al., used gene co-expression and protein interaction networks to predict 7 putative genes in breast cancer (Yue et al. 2016). The authors showed that the 7 gene signature could be a useful predictor of disease prognosis in breast cancer patients. A recent integrated network analysis and machine learning approach was able to accurately classify TNBC from non-TNBC based on a 16 gene signature (Naorem et al. 2019). Comprehensive transcriptomic and genomic analyses have identified novel molecular TNBC subtypes and subtype specific genes. Liu et al., (Liu et al. 2016) classified 165 TNBC tumors into four distinct subtypes, including immunomodulatory, luminal androgen receptor, mesenchymal-like, and basal-like and immune suppressed subtypes based on gene expression microarray data. Similar TNBC subtypes were identified by another study based on RNA and DNA profiling analyses of 198 tumors (Burstein et al. 2015). Although substantial research has been performed on breast cancer, including different molecular subtypes, there is a gap in understanding the molecular basis of each subtype. In addition, more focused studies are required to identify better therapeutic targets for the aggressive TNBC subtype.

In the current study, we have performed meta-analysis of microarray and RNA sequencing data to identify candidate genes associated with four molecular subtypes of breast cancer (luminal A, luminal B, HER2 and TNBC). Furthermore, protein interaction network analysis of the candidate genes was explored to identify functionally important molecules in each subtype. In addition, miRNA-gene interactions were explored to identify potential therapeutic targets for TNBC subtype.

## Methods

### Microarray and RNA sequencing data collection and processing

Literature search was performed using selected search engines, as indicated before (Bajpai et al., 2011). The microarray data corresponding to breast cancer molecular subtypes and breast normal tissues were downloaded from Gene Expression Omnibus (GEO) repository (Barrett et al. 2013). The raw data files of cancer and normal samples were pre-processed using *MAS5* algorithm (Hubbell et al. 2002) in a R-based environment to obtain absolute calls (‘P’, ‘A’, and ‘M’) of each gene in each sample. The normalized files with absolute calls were processed using a meta-analysis approach reported earlier (Acharya et al. 2010) to obtain genes with consistently transcribed status (‘P’) along with their reliability scores. Briefly, the method assigns a reliability score of +2 or −2 for a gene to be ‘transcribed’ or ‘dormant (A)’, respectively in each sample/hybridization. The score for each gene was then added across multiple samples/hybridizations to obtain a cumulative reliability score, in each subtype as well as normal breast tissue. The cumulative reliability score was then used to assign the final expression status (transcribed/dormant) for the genes. Genes with ‘transcribed’ status having a reliability score of at least 20 (present in at least 10 samples/hybridizations) were considered further. Differentially expressed genes in each breast cancer subtype compared to normal breast samples were obtained by using the GEO2R function implemented in GEO repository. The GEO2R function uses the normalized data submitted by the authors and applies Linear Models for Microarray data (‘*limma*’) (Ritchie et al. 2015) algorithm to identify differentially expressed genes between the groups. Genes with a │log2fold ≥2│ and FDR (B&H) adjusted *P* value of <0.01 in at least 2 independent studies were considered further.

The Cancer Genome Atlas (TCGA) RNA sequencing based raw read count data for the breast cancer molecular subtypes and normal breast samples were downloaded from Firebrowse cancer data portal (http://firebrowse.org/). Only primary tumor samples were considered for the analysis. Differentially expressed genes between each breast cancer molecular subtype and paired normal breast tissues were obtained using R based *edgeR* package (Robinson et al. 2010). Genes with │log2fold ≥2│ and FDR (B&H) *P* value of <0.01 were considered to be significant.

### Identification of candidate genes associated with breast cancer molecular subtypes and functional analysis

Meta-analysis of microarray and RNA-seq data was performed to identify candidate genes associated with breast cancer subtypes. Briefly, genes having both ‘transcribed’ and ‘up-regulated’ status in cancer (henceforth referred to as ‘up-regulated‘) and ‘transcribed in normal breast’ and ‘down-regulated’ status in cancer (henceforth referred to as ‘down-regulated’) based on microarray technology were considered as a preliminary-set of genes associated with each breast cancer subtype. Among these, genes in agreement with TCGA RNA-sequencing data were considered as the final set of candidate genes in each subtype. Based on the fold change obtained from TCGA-RNA-seq data, the candidate genes were classified at 3 levels of differential expression: low (log2 fold ≤3), moderate (log2 fold between ≥3.5 and ≤5 fold) or high (log2 fold ≥5.5).

The functional analysis, including Gene Ontology (GO), pathway, and chromosome enrichment analysis of the candidate genes was performed using DAVID (Huang da et al. 2009). Annotations with a *P*-value <0.05 were considered significant. The significant GO terms from each subtype were further submitted to REVIGO (Supek et al. 2011) to remove the redundancy and obtain a more meaningful set of annotations. Heatmaps of the commonly regulated candidate genes were constructed using the ClustVis tool (Metsalu and Vilo 2015) based on TPM values in each molecular subtype.

### Network analysis of the candidate genes associated with each breast cancer subtype

The known protein interactions (i.e., those with ‘experimental’ and ‘database’ evidence) among the differentially expressed candidate genes in each breast cancer subtype were obtained from the STRING database (Szklarczyk et al. 2019), as suggested to be suitable by a recent study (Bajpai et al., 2020) with a medium confidence threshold (confidence score, 0.4). The interactions were visualized and analyzed using the Cytoscape network analysis tool (Shannon et al. 2003), and MCODE tool (Bader and Hogue 2003) was used to identify protein clusters in each network. Significantly enriched (*P* <0.05) annotations for the selected clusters were obtained using the DAVID analysis tool (Huang da et al. 2009). Using CytoHubba (Chin et al. 2014), network statistics, such as node degree, betweenness and closeness centrality were determined. Significance analysis of the network topological features between different groups of differentially regulated genes (low, moderate and high) was performed using the two-tailed, unequal variance t-test. The miRNAs for the target genes were predicted using TargetScan (Agarwal et al. 2015), miRanda (Betel et al. 2008), and miRDB (Wong and Wang 2015) and the ones identified by all the three tools were considered further. The expression of miRNAs was verified using the literature as well as dbDEMC (Yang et al. 2017) and UALCAN (Chandrashekar et al. 2017) databases.

## Results

### Breast cancer and normal breast tissue samples used for derivation of candidate genes

A total of 8 microarray gene expression studies were identified from GEO repository corresponding to four different molecular subtypes of breast cancer (luminal A, luminal B, HER2, and TNBC). Among the eight studies identified, 6 had normal breast samples along with the cancer samples and were used for the derivation of differentially expressed genes. Two of the studies (GSE45827 and GSE31448) contained samples for all the four molecular subtypes. Triple-negative breast cancer had the highest number of samples (465) followed by luminal A (146), HER2 (95) and luminal B (83). The samples corresponding to only Affymetrix platforms were used for the derivation of absolute calls because of their compatibility with the meta-analysis approach used. Furthermore, additional normal breast samples collected from GEO (data not shown) were used for the derivation of absolute calls.

The TCGA RNA sequencing data was obtained for a total of 572 samples across four breast cancer molecular subtypes. Luminal A had highest number of samples (301) followed by luminal B (150), TNBC (95) and HER2 (26). There were 90 samples corresponding to normal breast tissue.

### Candidate genes associated with breast cancer molecular subtypes

Six microarray studies were used to identify differentially expressed genes in four breast cancer molecular subtypes compared to normal breast with a │log2 fold ≥ 2│ and FDR adjusted (B&H) *P* value of <0.01. Overall, the number of down-regulated genes obtained was more than that of the up-regulated genes in most studies. Triple-negative breast cancer (TNBC) subtype had the highest number of differentially expressed genes. Furthermore, 567 differentially expressed genes in luminal A, 458 in luminal B, 584 in HER2, and 720 genes in TNBC were reproduced by at least two microarray studies. Approximately, 80% of these genes also had ‘Transcribed’ status in breast cancer (for up-regulated genes) or normal breast samples (for down-regulated genes), and were considered as preliminary set of breast cancer-subtype associated candidate genes. These included 499 differentially expressed genes in luminal A, 411 in luminal B, 515 in HER2, and 638 genes in TNBC. RNA sequencing analysis identified a total of 10,938 genes to be differentially expressed across four breast cancer subtypes with a │log2 fold ≥2│ and FDR (B&H) *P*-value of <0.01. Among these, 5,766 genes were up-regulated and 5,172 were down-regulated. Triple negative breast cancer had maximum number of differentially expressed genes (2,950 genes).

A total of 1,487 candidate genes were identified based on the meta-analysis of microarray and RNA sequencing technologies (common to both the technologies) across four molecular subtypes of breast cancer. We identified 316 candidate genes (63 up- and 253 down-regulated) in luminal A, 347 in luminal B (86 up- and 261 down-regulated), 382 in HER2 (101 up- and 281 down-regulated), and 442 genes in TNBC (199 up- and 243 down-regulated) subtypes.

### Identification of COL10A1 as a potential diagnostic marker for breast cancer

The comparison of candidate genes among the subtypes showed 102 and 280 genes to be up- and down-regulated, respectively, in at least two molecular subtypes, whereas, 26 up- and 109 down-regulated genes were shared by all the four breast cancer subtypes (**Fig. 1**). In luminal A and luminal B subtypes, COL10A1 had the highest fold change (7.6-fold, *P* = 2.19E-89 and 7.4-fold, *P* = 5.74E-63, respectively), whereas, in HER2 and TNBC subtypes, MMP1 had the highest fold change for up-regulation (8.9-fold, *P* = 1.23E-35 and 8.2-fold, *P* = 2.44E-42). Among the top 10 up-regulated genes (based on fold change), COL10A1, COL11A1, and MMP11 were common across all four subtypes. While, COL10A1 and COL11A1 were up-regulated with at least 6 fold, MMP11 was up-regulated with more than 5.5 fold in all the subtypes. When the average fold was considered across all the four subtypes, COL10A1 had a highest mean fold of 7.1, followed by COL11A1 and MMP11 with 6.8 and 6.3 folds, respectively. In addition, the mRNA and protein expression of COL10A1 was confirmed to be significantly higher in breast cancer molecular subtypes compared to normal breast tissues (Chandrashekar et al. 2017). Thus, COL10A1 may be used as a potential diagnostic marker for breast cancer, irrespective of the subtype, whereas RAB26 and MMP1 may be specifically used as markers for ER/PR-positive and - negative breast cancers, respectively. Among the down-regulated genes, LEP and CA4 were in the top 5 in all four subtypes. Survival analysis indicated that among the commonly up-regulated genes, KIF4A was significantly associated with the survival of breast cancer patients, whereas, among the commonly down-regulated genes, significant association was observed for ADIPOQ, C2orf40, CD300LG, CIDEA, CLDN11, CNN1, and SDPR.

**Fig. 1.**
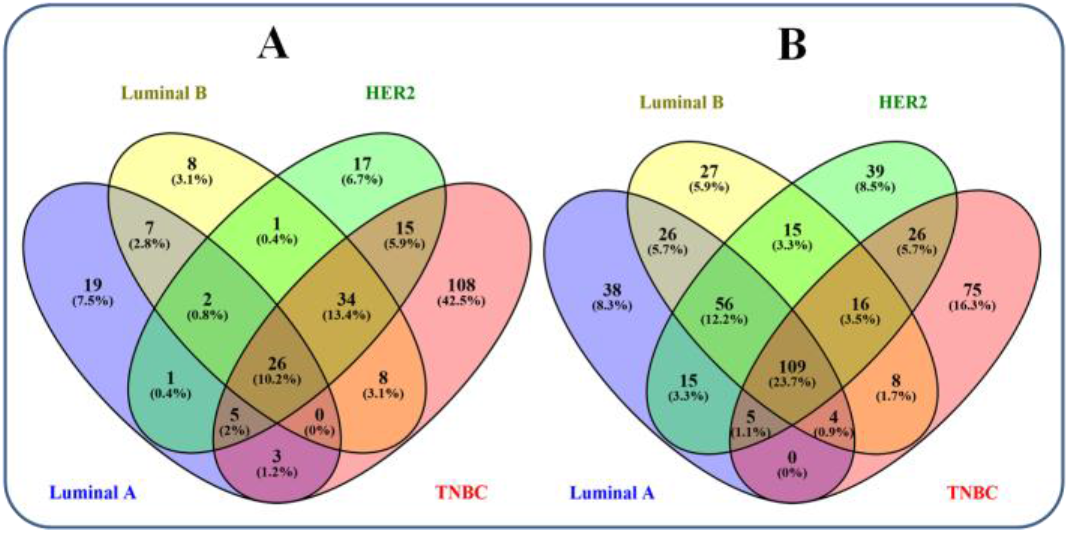
Candidate genes shared across four breast cancer subtypes. A: up-regulated genes; B: down-regulated genes.

### Functional analysis of the candidate genes associated with breast cancer molecular subtypes

The functional analysis of the candidate genes in each breast cancer subtype showed enrichment of multiple processes and pathways. Processes, such as cell division, cell proliferation, apoptosis, and chromosome segregation were found to be significantly enriched by the up-regulated genes in all the breast cancer subtypes. The down-regulated genes were significantly enriched for metabolic related processes, including cholesterol homeostasis, glucose homeostasis, and lipid metabolic process. Pathway enrichment analysis showed up-regulated genes to be significantly enriched for ‘cell cycle’ and ‘p53 signaling’ pathways in all four molecular subtypes (**Fig. 2**). The up-regulated genes in luminal A were specifically enriched for ‘PI3K –Akt signaling’, ‘focal adhesion’, and ‘protein digestion and absorption’ pathways. The up-regulated genes in TNBC were enriched for multiple pathways including, ‘Fanconi anemia’, ‘HTLV-I infection’, and ‘homologous recombination’ pathways. The down-regulated genes in each subtype were significantly enriched for multiple signaling pathways, such as ‘PPAR’, ‘adipocyte’, and ‘MAPK’ signaling pathways. Other commonly down-regulated pathways in all four subtypes include ‘focal adhesion’, ‘Tyrosine metabolism’, ‘regulation of lipolysis’, and ‘drug metabolism-cytochrome P450’. Many pathways were specifically down-regulated in each breast cancer subtype. For example, ‘glycolysis’ and ‘pyruvate metabolism’ in luminal A, ‘Leukocyte transendothelial migration’, in luminal B, multiple signaling pathways in HER2, and ‘ABC transporter’, ‘nitrogen metabolism’, ‘renin secretion’, and ‘axon guidance’ in TNBC subtype (**Fig. 2**). Analysis of 26 commonly up-regulated genes showed ‘cell cycle’, ‘p53 signaling,’ and ‘oocyte meiosis’ pathways to be significantly enriched. The 109 commonly down-regulated genes were significantly enriched for multiple signaling pathways.

**Fig. 2.**
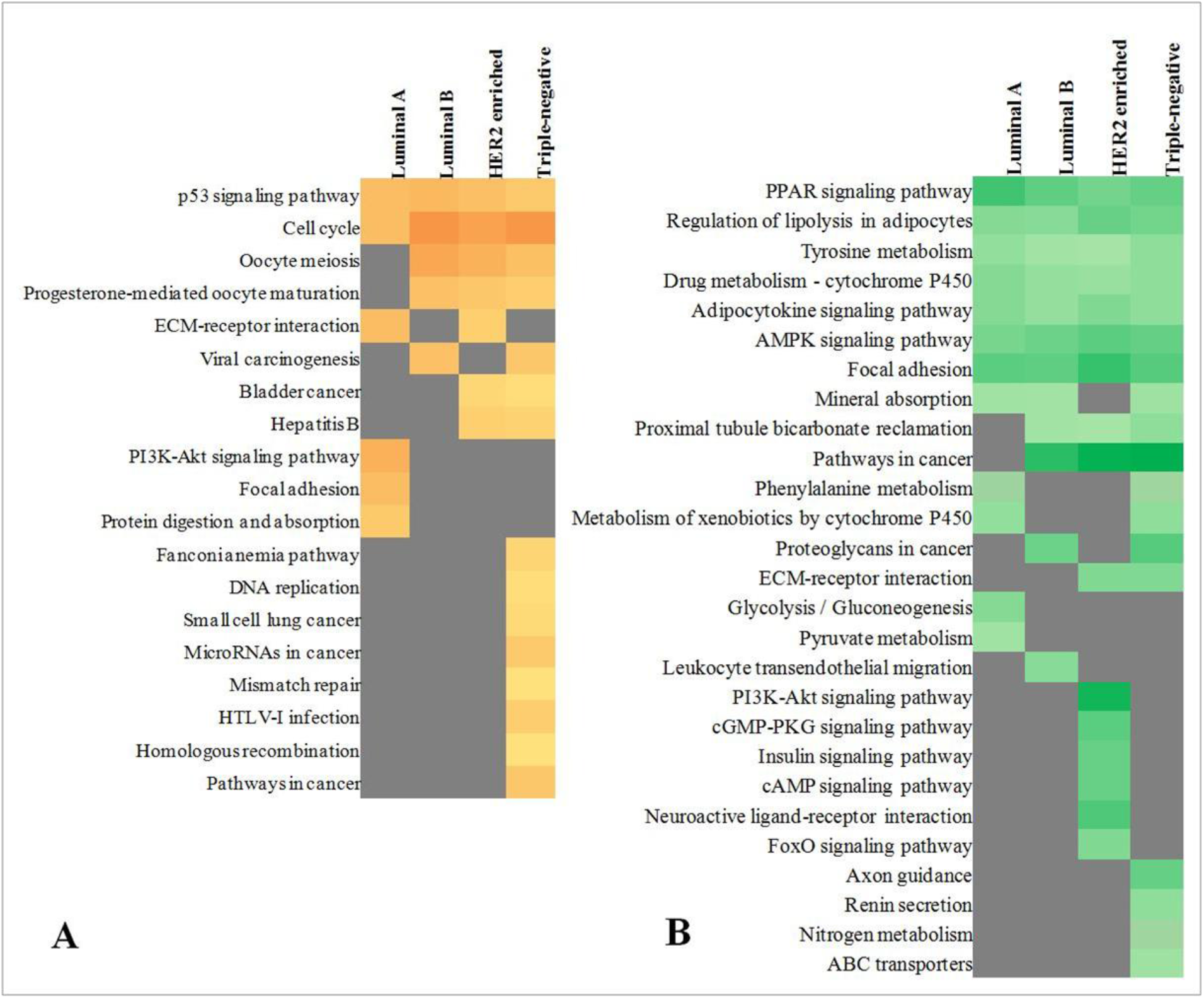
KEGG pathways significantly enriched (*p* <0.05) by (A) up-regulated and (B) down-regulated genes in breast cancer subtypes. Orange/Green box: presence of pathway; dark grey box: absence of pathway. The percentage of genes involved in each pathway in each subtype is represented by the intensity of orange/green color (dark color: higher percentage; light color: lower percentage).

Chromosomal distribution showed significant enrichment of chromosome 1, 4, 8, 15, 17, and 20 for the up-regulated candidate genes in different molecular subtypes. While chromosome 15 and 17 were most significantly enriched for the up-regulated genes in luminal B (*P* = 0.014) and HER2 (*P* = 0.015) subtypes, respectively, chromosome 4 and 20 were most significantly enriched for luminal A (*P* = 0.006) and TNBC (*P* = 0.022) subtypes, respectively. For the down-regulated genes, chromosome 3 was most significantly enriched in three breast cancer subtypes (luminal B, *P* = 0.018; HER2, *P* = 0.0009; TNBC, *P* = 0.028) covering around 10% of the genes in each, whereas, chromosome 11 was most significantly enriched for the down-regulated candidate genes in luminal A (*P* = 0.029) subtype (covering ~10% of the genes). As expected, there were no genes from Y chromosome in any subtype.

### Protein interaction network analysis of candidate genes associated with breast cancer molecular subtypes

Protein-protein interaction network was constructed among the differentially expressed candidate genes for each breast cancer subtype. The up-regulated genes were found to be more tightly connected than the down-regulated genes. Overall, highly up- or down-regulated genes had comparatively lesser number of interactions. The protein interaction networks along with the analysis of top 3 clusters in each subtype have been described below.

The protein interaction network revealed that 143 candidate genes in luminal A were connected with 220 edges (**Fig. 3**). Cluster analysis of the network identified a total of 10 clusters containing 42 proteins, of which 16 proteins (38%) were covered in the top 3 clusters (**Fig. 3**). The cluster 1 was most significantly enriched for ‘chemokine signaling’ and ‘cytokine-cytokine receptor interaction’ pathways, whereas, cluster 2 represented processes, such as ‘muscle contraction and ‘actin filament organization’, and cluster 3 was significantly enriched for ‘cell cycle’, and ‘p53 signaling’ pathways (**Fig. 3**). Furthermore, analysis of the protein interaction network using identified EGFR to be the key hub molecule, followed by PPARγ and CDK1 based on betweenness centrality. EGFR and PPARγ were down-regulated, whereas, CDK1 was up-regulated in luminal A subtype.

**Fig. 3.**
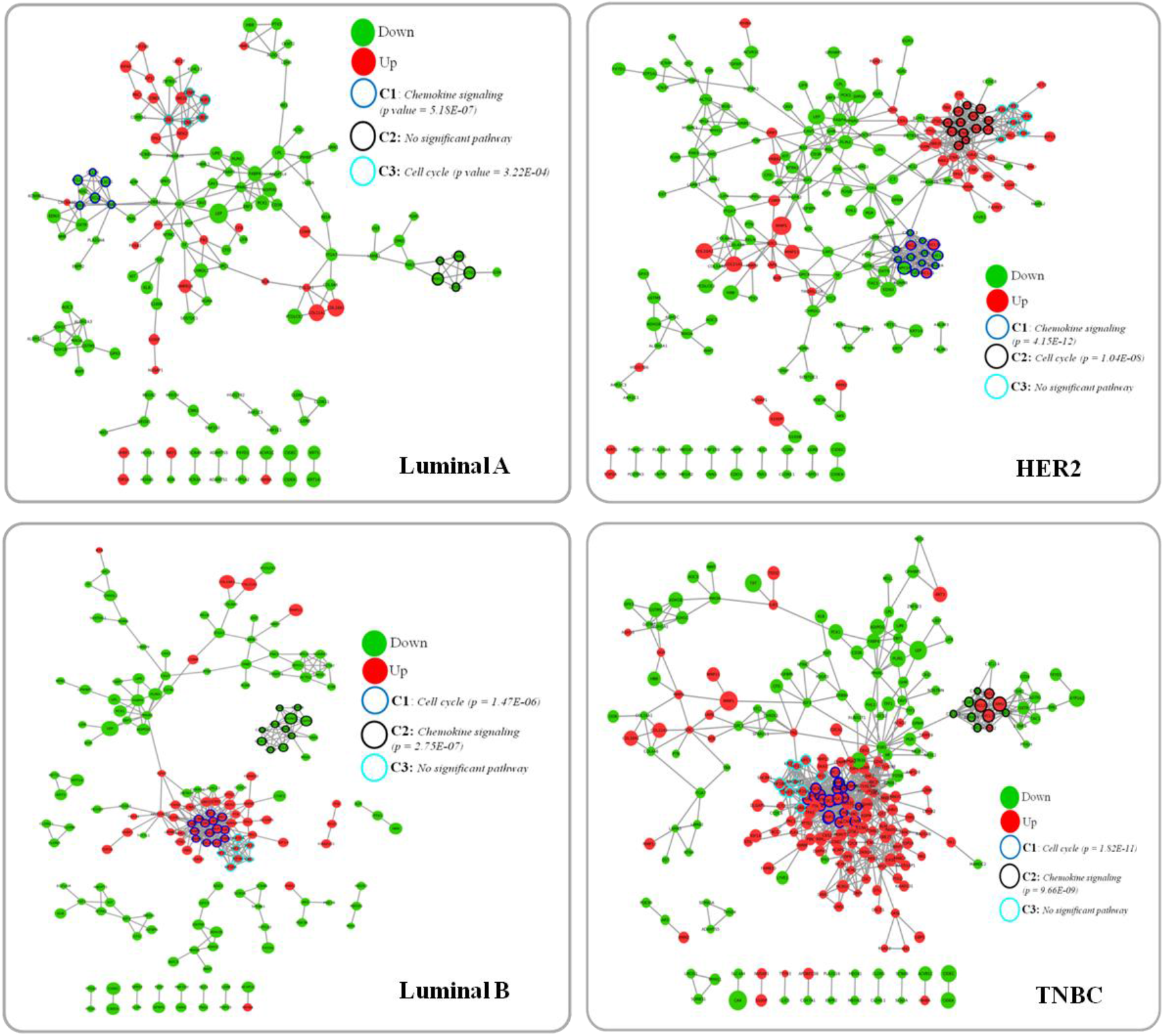
Protein interaction network of candidate genes in breast cancer subtypes. C1: Cluster1, C2: Cluster 2, C3: Cluster 3. Node size indicates the increasing fold change. Most significantly enriched pathway for each cluster is mentioned.

Among the 347 candidate genes in luminal B, 172 were found to be connected with 393 edges (**Fig. 3**). Cluster analysis of the protein interaction network identified 12 clusters with 64 nodes, 50% of which were covered by the top 3 clusters (**Fig. 3)**. Cluster 1 was highly connected (91 edges) with 14 up-regulated nodes and was significantly enriched for cell cycle related processes and pathways. Based on functional annotations, genes in cluster 1 of luminal B and cluster 3 of luminal A were involved in similar processes and pathways. Cluster 2 had 11 down-regulated genes and was significantly enriched for ‘chemokine signaling’. Cluster 3 contained 7 up-regulated genes connected with 21 edges, and was enriched for processes, such as ‘microtubule based movement’, ‘antigen processing and presentation’, ‘vesicle mediated transport’, and ‘cytokinesis’. Based on the betweenness centrality, PPARγ was found to be the key node, followed by EZH2 and CAV1. PPARγ and CAV1 were down-regulated, whereas, EZH2 was up-regulated in luminal B subtype.

Among the 382 differentially expressed candidate genes in HER2 subtype, 212 genes were connected with 542 edges (**Fig. 3**). The analysis of the protein interaction network identified 13 clusters with 83 nodes, 33 of which (15%) were represented by the top 3 clusters (**Fig. 3**). Cluster 1 having 14 genes was highly connected with 91 edges and was most significantly enriched for ‘chemokine signaling pathway’. Cluster 2 containing 13 up-regulated genes was significantly enriched for ‘cell cycle’ and 3 other pathways. Cluster 3 having 6 up-regulated genes was enriched for processes related to ‘microtubule-based movement’, ‘cell cycle’ and ‘antigen processing and presentation’. ESR1, a down-regulated gene in HER2 subtype, was identified as the key node based on betweenness centrality measure. Among the up-regulated genes, CDK1 and SDC1 were among the top ten hub molecules.

A total of 256 genes were found to be connected with 973 edges in triple-negative breast cancer protein interaction network (**Fig. 3**). The cluster analysis of the network resulted in a total of 18 clusters, highest among all the subtypes. Cluster 1 contained 23 up-regulated highly connected (253 edges) genes and was significantly enriched for ‘cell cycle’ and 5 other pathways. Custer 2 contained 5 up-regulated and 6 down-regulated genes and was significantly enriched for ‘chemokine signaling’, ‘cytokine-cytokine receptor interaction’, and ‘Toll-like receptor signaling’ pathways. Cluster 3 contained 10 up-regulated genes, all of which were associated with ‘microtubule-based movement’. ESR1, a down-regulated gene with a fold change of 4.2 was identified as the top hub gene in TNBC subtype, whereas, among the up-regulated genes, FN1 (fibronectin-1) was the key hub molecule with highest betweenness centrality measure.

### Identification of functionally important genes in breast cancer subtypes

The protein interaction network analysis across breast cancer subtypes identified functionally important molecules based on multiple topological features (**Fig. 4**). The results show that the average node degree of the low (log2 fold ≤3) and moderately differentially regulated genes (log2 fold between ≥3.5 & ≤5) was significantly higher compared to that of highly differentially regulated genes (log2 fold ≥5.5) in all breast cancer subtypes, except in luminal A (**Fig. 4A**). Similar trend was observed, when the percentage of nodes with a degree of at least 5 were compared. In case of luminal B, HER2 and TNBC subtypes, 38, 48, and 58% of the moderately differentially regulated genes, respectively were found to be interacting with at least 5 proteins, whereas, <25% of the highly differentially regulated genes in these subtypes had a degree of at least 5 (**Fig. 4B**). When global network topological features, such as betweenness and closeness centrality were explored, overall, highly differentially regulated genes had significantly lower score compared to the low and moderately differentially regulated genes (**Fig. 4C and D**). For example, EGFR in luminal A, PPARγ in luminal B, and ESR1 in HER2 and TNBC subtypes had a fold change of less than 5, but had highest betweenness centrality score. However, COL10A1, which had a very high fold change (> 6) in all the breast cancer subtypes, interacted with only 4 genes in any subtype and its betweenness centrality score was zero. Similarly, MMP11 was up-regulated with a fold change of more than 5.5 in all the four subtypes; however, it interacted with only 1 gene in three subtypes, and had no interaction in luminal A subtype. LEP, a highly down-regulated gene in all the four subtypes (≥5.8 fold) interacted with a maximum of 6 proteins and had a low betweenness centrality score. These findings suggest that the genes differentially expressed with a low-moderate fold change play an important role in the information flow across the proteins in breast cancer subtypes and may be potential therapeutic targets.

**Fig. 4.**
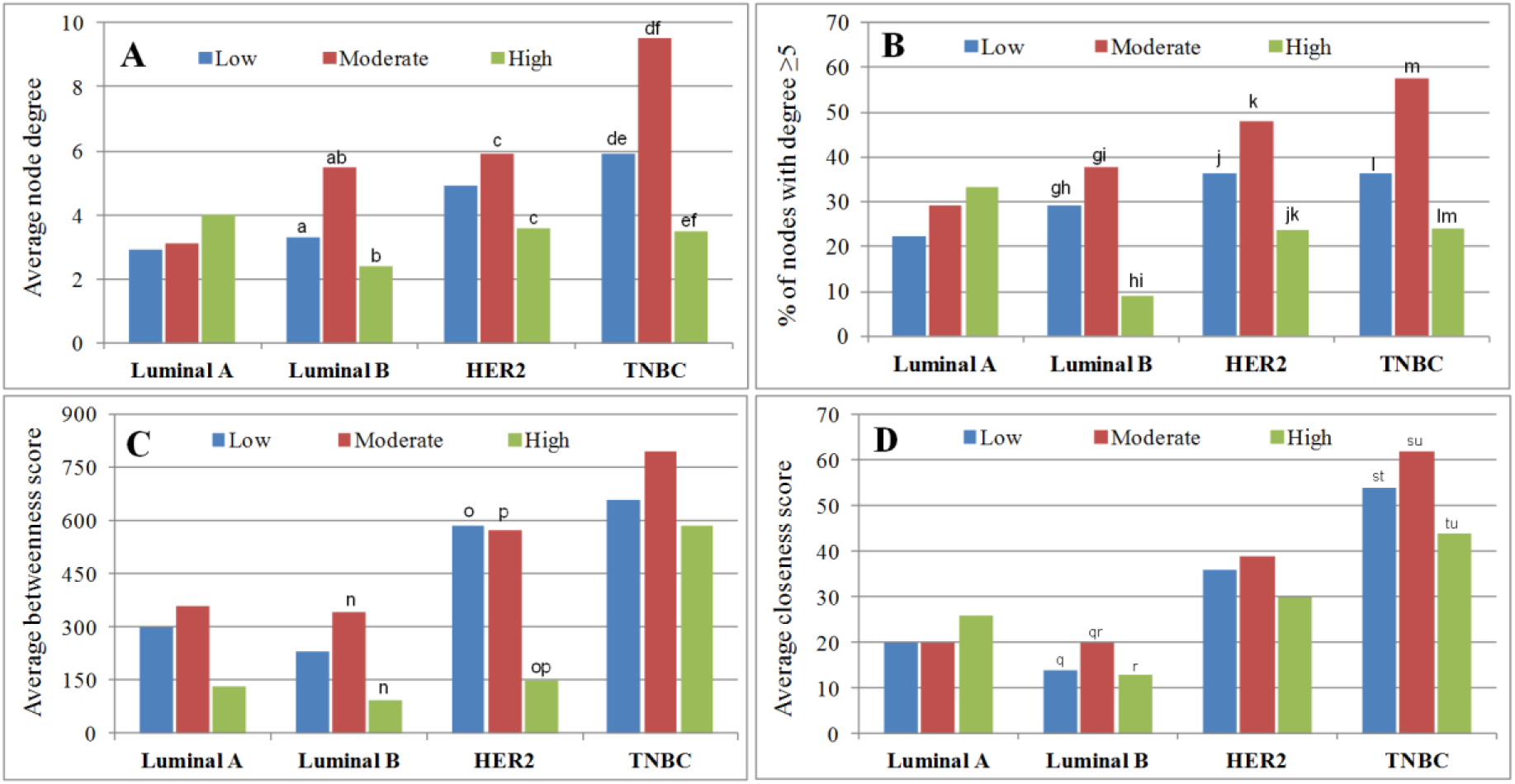
Topological features of the protein interaction network across breast cancer subtypes for low (log2fold ≤3), moderate (log2fold between ≥3.5 & ≤5), and high (log2fold ≥5.5) deregulated genes. A & B: Node degree; C: Betweenness centrality; D: Closeness centrality. The bars sharing same alphabetical letter are significantly different from each other with a p value <0.05. Only significant values are indicated. **a:** 0.0096; **b:** 0.0003; **c:** 0.0193; **d:** 0.0077; **e:** 0.0219; **f:** 4.2E-06; **g:** 0.0043; **h:** 0.0002; **i:** 1.5E-06; **j:** 0.0001; **k:** 2.5E-05; **l:** 0.0298; **m:** 0.0029; **n:** 0.0280; **o:** 0.0029; **p:** 0.0095; **q:** 0.0195; **r:** 0.0188; **s:** 0.0307; **t:** 0.0493; **u:** 0.0006.

### Interaction between FN1 and miR-1271-5p may be explored as a potential therapeutic option for TNBC

The FN1 gene was found to be up-regulated in TNBC subtype with a log 2 fold of 2.5, and had the highest betweenness among the up-regulated genes in TNBC. Hence, we explored its therapeutic potential using mRNA-miRNA interactions. Thirteen miRNAs were predicted to target FN1 gene based on three different methods (TargetScan, miRanda, and miRDB) (**Fig. 5**). Of these, miR-200c-3p, miR-200b-3p, and miR-1-3p were also found to be reported in miRTarBase (Chou et al. 2018), which stores experimentally proven interactions. Furthermore, among the 13 miRNAs, 12 have been reported in context of TNBC subtype in literature. However, there was no literature evidence indicating a direct association of hsa-miR-1271-5p with TNBC, although its association with breast cancer has been reported in few studies (Du and Liu 2018; Liu et al. 2019). Additionally, hsa-miR-1271-5p is down-regulated in breast cancer tissues compared to normal breast tissues based on the evidences from literature (Du and Liu 2018; Liu et al. 2019) as well as miRNA expression databases. Thus, miR-1271-5p mimics can be explored as a potential therapeutic option for TNBC subtype.

**Fig. 5.**
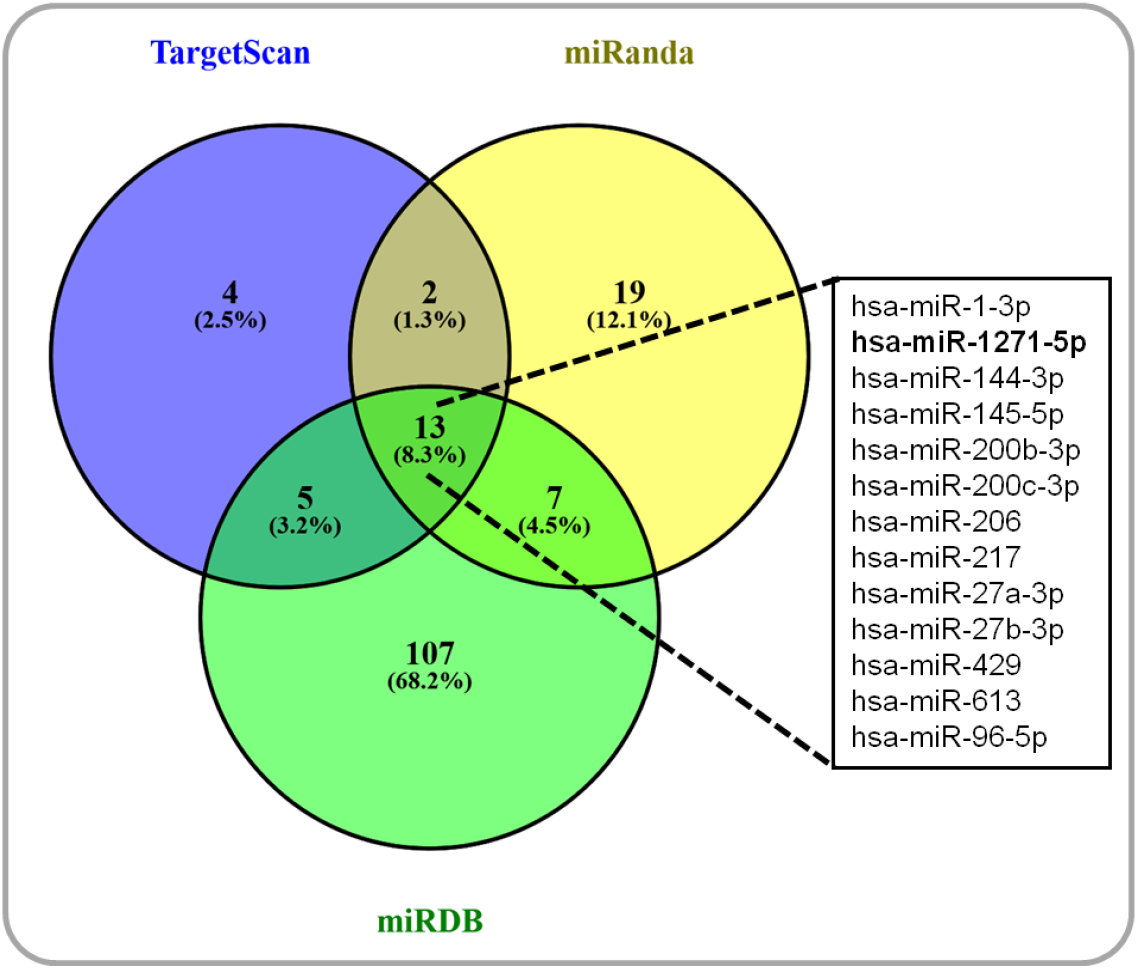
miRNAs targeting FN1 gene. The box provides a list of miRNAs targeting FN1 gene, common to all the three prediction methods (TargetScan, miRanda and miRDB). Novel miRNA targeting FN1 gene in TNBC subtype is highlighted in bold.

## Discussion

Meta-analysis of transcriptomic data is known to provide novel insights into the molecular details of cancer (Davuluri et al., 2021). In the current study, we have performed meta-analysis of publicly available microarray and RNA-sequencing data across four major molecular subtypes of breast cancer (luminal A, luminal B, HER2 and TNBC). Although RNA sequencing data contributed comparatively higher number of genes; microarray studies provided higher sample size, particularly for HER2 and TNBC subtypes. We identified candidate genes strongly associated with each molecular subtype as well as genes commonly differentially expressed in all four subtypes. The functional analysis of the candidate genes in each subtype showed up-regulation of ‘cell cycle’ and ‘p53 signaling’ pathways, corroborating with findings from other studies involving breast and other cancers (Peng et al. 2017; Tian et al. 2018). Furthermore, the current study showed down-regulation of multiple signaling pathways (PPAR, AMPK, and adipocytokine signaling) in breast cancer subtypes. AMPK signaling has been reported to prompt cells to produce energy at the expense of growth and motility (Zhao et al. 2017). Hence, decreasing AMPK activity may induce proliferation and metastasis of tumor cells. Multiple studies have reported that activation of AMPK signaling leading to apoptosis of breast cancer cells (Lin et al. 2010; Lin et al. 2017; Song et al. 2017).

Furthermore, enrichment analysis showed chromosome 1 to be significantly enriched by the up-regulated genes in two subtypes, and chromosome 3 by the down-regulated genes in three subtypes. Study by Soloviev et al. (Soloviev et al. 2013) has reported significant overexpression of 26 genes mapped to chromosome 1q in breast cancer compared to normal breast tissues. Another study (Orsetti et al. 2006) identified 30 candidate genes from chromosome 1 showing significant overexpression correlated to copy number increase using microarray. Multiple genes mapping to chromosome 3 display low expression in breast cancer tissues, generally due to the loss of chromosome 3 regions (Al Sarakbi et al. 2009; Silva et al. 2006; Thompson et al. 2014; Zia et al. 2007).

The genes, MMP11, COL11A1, and COL10A1 were up-regulated in four breast cancer subtypes with a log 2 fold value of >5.5. The role of MMP11 has been well established in multiple cancers including breast cancer (Pang et al. 2016; Zhang et al. 2016). Few studies have also shown that MMP11 can be used as a biomarker for metastatic breast cancer (Gonzalez de Vega et al. 2019; Roscilli et al. 2014). Similarly, COL11A1 has also been studied as a marker in breast cancer (Freire et al. 2014) as well as other cancers (Garcia-Pravia et al. 2013; Wu et al. 2014). The role of COL10A1 has been explored in multiple cancers (Chapman et al. 2012) and recently in breast cancer subtypes (Zhang et al. 2020). This gene encodes an alpha chain of type X collagen and it is a component of extracellular matrix. Mutations in COL10A1 are associated with Schmid type metaphyseal chondrodysplasia (Warman et al. 1993) and Japanese type spondylometaphyseal dysplasia (Ikegawa et al. 1998). Chapman et al., (Chapman et al. 2012) reported elevated expression of COL10A1 in diverse tumor types and restricted expression in human normal tissues. The authors stated that COL10A1 can be used as a novel target for diagnosis and treatment of multiple cancer types. In addition, increased expression of stromal COL10A1 correlates with poor pathological response in ER+/HER2+ breast tumors (Brodsky et al. 2016). Nevertheless, the current study shows that COL10A1 is highly up-regulated in all breast cancer subtypes; therefore, it may be used as a diagnostic marker for breast cancer, irrespective of the subtype.

The degree and betweenness are important measures for identifying functionally important genes/proteins in a protein interaction network (Yu et al. 2007). While degree, which indicates the number of connected nodes, is a local network property; betweenness centrality is a global network property and indicates the nodes important for information flow across the complete network. Joy et al. (Joy et al. 2005), found that the proteins with high betweenness score are more likely to be essential in a yeast protein interaction network. Further, a mammalian transcription network clearly displays a high degree and betweenness score for the tumor-suppressor and proto-oncogenes (Potapov et al. 2005). In the current study, we showed that low-moderately differentially expressed genes had high betweenness score and may be important for information flow across the protein interaction network in breast cancer subtypes than the highly-deregulated genes. Hence, they can be better therapeutic targets.

Fibronectin 1(FN1) gene was found to be the key up-regulated ‘hub gene’ in TNBC subtype based on the highest betweenness centrality score with a fold change of 2.5. It encodes a fibronectin protein, which is found on the cell surface and in extracellular matrix and is known to be involved in cell adhesion, motility, and wound healing (Li et al. 2017b; Xiao et al. 2018; Yoshimura et al. 2018). Being central to the protein interaction network, suppression of FN1 may disrupt the information flow across multiple proteins, which may affect multiple processes and pathways, and hence targeting FN1 may help in tumor inhibition. In addition, its involvement in cell adhesion and migration process makes it a potential candidate drug target. Recent studies have also shown its association with multiple neoplasms. High stromal expression of FN1 has been shown to be associated with aggressive pancreatic carcinoma (Hu et al. 2019), although the study did not find any association with overall survival of patients. A study by Yoshimura et al. (Yoshimura et al. 2018) showed that miR-99a-5p promotes cell invasion in ovarian cancer by up-regulating fibronectin and vitronectin. Another study (Xiao et al. 2018) showed that high FN1 expression in esophageal carcinoma is closely associated with poor prognosis. The high stromal content of the protein facilitates tumor cell metastasis by promoting morphological changes and improving the motility of cancer cells. Other cancers where high FN1 expression has been reported include lung adenocarcinoma (Li et al. 2017b), prostate cancer (Petrini et al. 2017), and hepatocellular carcinoma (Liu et al. 2017). The role of FN1 has also been reported in breast cancer detection, development, and metastasis (Hong et al. 2014; Jeon et al. 2015; Moon et al. 2016), including TNBC and other breast cancer subtypes (Suman et al. 2016). Study by Wang et al., (Wang et al. 2011) showed that miR-1 suppresses migration and invasion of laryngeal carcinoma by negatively regulating FN1 gene. In the current study, we identified FN1 gene to be targeted by miR-1271-5p based on three different prediction methods. Multiple studies have reported the anti-cancer role of miR-1271 in various cancers (Li et al. 2017a; Wang et al. 2018; Xie et al. 2018). In addition, a few studies have reported its tumor suppressor activity in breast cancer via targeting SNAI2 (Liu et al. 2019) and SPIN1 (Du and Liu 2018) genes. However, the relationship between miR-1271-5p and FN1 has not been reported in breast cancer till date, although their interaction has been experimentally validated in glioma (Gong et al. 2017). Our results indicate that the interaction between FN1 and miR-1271-5p can be exploited as a potential therapeutic option for TNBC. However, *in vitro* and *in vivo* studies are required to confirm the role of miR-1271-5p in the suppression of FN1 gene in TNBC subtype.

The current study had a few limitations. While categorizing the genes into different regulation-based groups (low, moderate and high), the fold change threshold was considered based on the minimum and maximum fold change values of the candidate genes in breast cancer subtypes. However, the fold change threshold for deriving such genes may vary depending on the condition of interest. For instance, in the current study, genes with │log2fold <3│ were considered as lowly deregulated. However, same criteria may not hold true for other cancer types or conditions, where the fold difference between the test and control conditions is not considerably high. Furthermore, the functional importance of the low-moderately deregulated genes in breast cancer subtypes need to be confirmed experimentally.

## Conclusions

Meta-analysis of microarray and RNA sequencing data identified candidate genes associated with four major subtypes of breast cancer. Functional analysis of these genes showed up-regulation of ‘cell cycle’ and ‘p53 signaling’ pathways, and down-regulation of ‘PPAR’, ‘Adipocytokine’ and ‘AMPK’ signaling, ‘drug metabolism’, ‘tyrosine metabolism’, ‘regulation of lipolysis’ and ‘focal adhesion’ pathways in all four subtypes of breast cancer. While multiple chromosomes were found to be significantly enriched by the up-regulated genes, chromosome 3 had a significantly higher share for the down-regulated genes in majority of the subtypes. COL10A1 was found to be significantly up-regulated in all breast cancer subtypes with a high fold (mean log 2 fold of 7.1) and may be used as a potential diagnostic marker for breast cancer, irrespective of the molecular subtype. Among the commonly up-regulated candidate genes, the analysis showed an association of KIF4A with the survival of breast cancer patients. Significant association was observed for seven of the commonly down-regulated candidate genes. Protein interaction network analysis of the candidate genes identified distinct clusters of proteins and hub molecules in each subtype. Moreover, the network analysis showed that low-moderately deregulated genes control the information flow across proteins in breast cancer subtypes, and hence may be better therapeutic targets. MiR-1271-5p may have a role in the negative regulation of FN1 gene, an up-regulated and functionally important hub gene in TNBC. Hence, synthetic miR-1271 mimics may be used as a potential therapeutic option for treating triple-negative breast cancer. However, experimental validations are required to confirm the regulatory role of miR-1271-5p in TNBC.

## Acknowledgements

This research did not receive any specific grant from funding agencies in the public, commercial, or not-for-profit sectors. Research in IBAB is supported by the IT, BT & ST department of the Government of Karnataka.

## Author contributions

**AKB:** Conceptualization, Methodology, Data curation, Validation, Formal analysis, Investigation, Visualization, Writing - Original Draft; **SD:** Data curation, Validation, Visualization, Investigation, Writing - Original Draft; **KT:** Supervision, Writing - Review & Editing; **KKA:** Conceptualization, Methodology, Supervision, Project administration, Writing - Review & Editing.

